# Convincing Evidence of Carnivorous Diet in Alvarezsaurian Dinosaur

**DOI:** 10.1101/2024.12.06.627300

**Authors:** Shuo Wang, Nuo Ding, Waisum Ma, Wenmiao Yu, Tingting Zheng, Jonah Choiniere, Xing Xu

## Abstract

A dietary shift from carnivory to insectivore has been proposed to explain the dramatic morphological evolution of alvarezsaurians, particularly the adaptation related to the manual digital reduction and body size miniaturization. However, based solely on morphological shifts, this hypothesis lacked direct dietary evidence to support either carnivory or insectivore. Here, we present the first convincing dietary evidence for alvarezsaurians, derived from the intestinal contents of the Early Cretaceous *Bannykus wulatensis*. Our analysis revealed significantly higher levels of calcium and phosphorus in the intestinal contents compared to the surrounding sandstone. Scanning electron microscopy identified hard tissue debris and possible soft tissues surrounding by phosphatized bacteria and tightly packed hollow microspheres, suggesting that the intestinal contents were strongly pseudomorphed by phosphatized microbes during fossilization. Raman spectroscopy showed characteristic peaks indicative of bone-derived material, consistent with the hard tissue debris appeared in the intestinal contents. Our results suggest that *Bannykus* had a carnivorous diet with strong chemical digestion, which likely compensated for its delicate cranial structures and small teeth. These results imply that if a dietary shift to insectivore occurred, it likely took place later in alvarezsaurian evolution, probably coinciding with a reduction in body size.

---

Reconstructing the diet and possible feeding habits of extinct animals is one of the major tasks for modern paleobiology[1]. While direct indicators such as stomach and intestinal contents[2–4], gastric pellet[5, 6], and coprolites[7–9], can reveal diets, these are rarely preserved unless non-digestive hard tissues including bone fragments[2–4, 9], seed coats[10, 11]or plant fragments[7] are present[11–13]. This makes carnivores[3] (including osteophagus[8, 9] and piscivorous[5]), granivores[10] and frugivores[14, 15] easier to be identified in the fossil record. In contrast, insectivores, folivores, nectarivores, saprophages, myrmecophagies, omnivores and detritivores are typically inferred from indirect evidence, such as isotopic signals[16], comparative functional morphological of teeth and jaws[14], dental microwears[17], and the presence or absence of gastroliths[1].

Alvarezsaurians are a group of small, lightly-built maniraptoran dinosaurs with seemingly conflicting characteristics: their elongated hindlimbs suggest strong cursorial abilities, while the absence of gastroliths implies they were unlikely to have developed a gastric mill. However, their delicate cranial elements, presence of cranial kinesis, tiny and tightly-packed simplified homogenous teeth indicating limited bite force, and the highly reduced manual digits in later-diverging taxa[18, 19] make their feeding strategies controversial. Some studies proposed that the monodactyl hands of later-diverging alvarezsaurids were specifically adapted for digging into ant nest dunes (myrmecophagous)[19–21], but others argued that the rather limited movement of these manual elements makes digging or burrowing actions unlikely[22–24]. In contrast, the earliest known alvarezsauroid, *Haplocheirus*, possesses unreduced grasping manual digits and larger, recurved serrated teeth[25], indicating it may have occupied a different ecological niche compared to their later-diverging relatives. While a dietary transition has been hypothesized to have accompanied the dramatic morphological changes in alvarezsaurian evolution[26], direct dietary evidence supporting either carnivory or insectivore including myrmecophagous has been lacking. This makes the search for such evidence crucial to testing this hypothesis [25, 26].

Here, we report that *Bannykus wulatensis* (IVPP V25026)[27], an Early Cretaceous alvarezsauian that is temporally, morphologically, and functionally intermediate between *Haplocheirus* and later-diverging alvarezsaurians, was a carnivore base on the analysis of its intestinal contents. Further analysis suggests that *Bannykus* might have developed a highly efficient chemical digestion to break down the bone components in its food, which is unique among known theropods.

## Results

Further preparation of the type specimen of *Bannykus wulatensis* revealed a three-dimensionally preserved posterior gastralia ribcage, indicating minimal mechanical disturbance to the abdominal region since death. A yellowish conglomerate beneath the last dorsal vertebrae, corresponding to the duodenum-to-rectum region seen in *Scipionyx samniticus*[2] and *Sinocalliopteryx gigas*[28], is enclosed by the gastralia (Fig. 1). The distinct color and heterogeneous texture of this conglomerate compared to the surrounding greyish-green sandstone (Fig. 1, b-d) suggest that some abdominal contents, including food remains, may have been preserved *in situ*.

**Figure 1.**
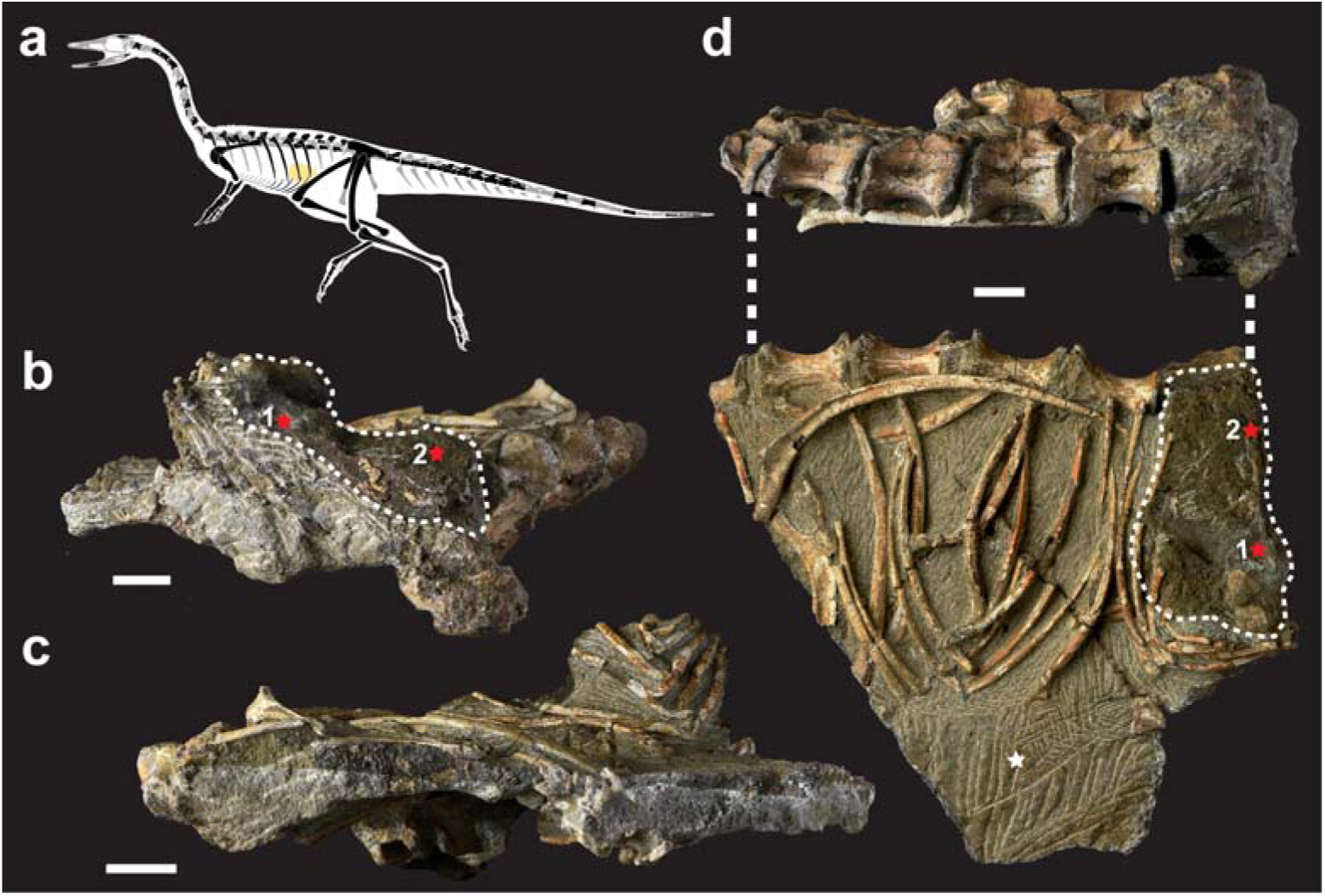
Silhouette and photographs of *Bannykus wulatensis* holotype (IVPP V25026) showing the yellowish conglomerate enclosed by the partial gastralia ribcage. **a,** Skeletal silhouette of *Bannykus wulatensis* indicating the relative position of the yellowish conglomerate enclosed by the gastralia ribcage (not to scale); **b-d,** Close-up images of the block containing the yellowish conglomerate in posterior view (**b**), anterior view (**c**), left lateral view (**d, upper**), and ventral view (**d, lower**). Notably, the three-dimensionally preserved partial gastralia ribcage is visible in **c**. The white dashed line in panels **b** and **d** outlines the yellowish conglomerate within the ribcage, while the red and white stars in panels **b** and **d** marks the positions where samples of conglomerate and host sandstone were taken, respectively; Arabic numerals indicate the sample numbers. **Scale bar**=2cm.

To test this speculation, two samples were taken from the conglomerate (Fig. 1b, d, red stars), along with a control sample from sandstone outside the ribcage (Fig. 1d, white star). Thin sections revealed that the control sample is calcareous-cemented sandstone, whereas the abdominal conglomerate samples consist of angular mineral particles embedded in a fine-crystalized matrix (Figs. 2-3). The mineral particles that make up the sandstone are more rounded, indicating they had been transported over longer distances compared to those in the abdominal conglomerate. Energy-dispersive spectroscopy (EDS) and electron microprobe (EMP) analysis revealed significantly higher levels of calcium and phosphorus in the conglomerate (Figs. 2-3, Extended Data Table 1), primarily in the forms of apatite (with the phosphorus elemental map serving as a proxy for calcium phosphate[6, 29]). This result is consistent with findings from other bone-rich intestinal contents[2, 6, 29] or coprolites [8, 29]. In contrast, the host sandstone contains heavy minerals such as ilmenite and hematite (Figs. 2-3, Extended Data Table 1). These lines of evidence suggest the apatite were originated from *Bannykus* remains rather than host sediments.

**Figure 2.**
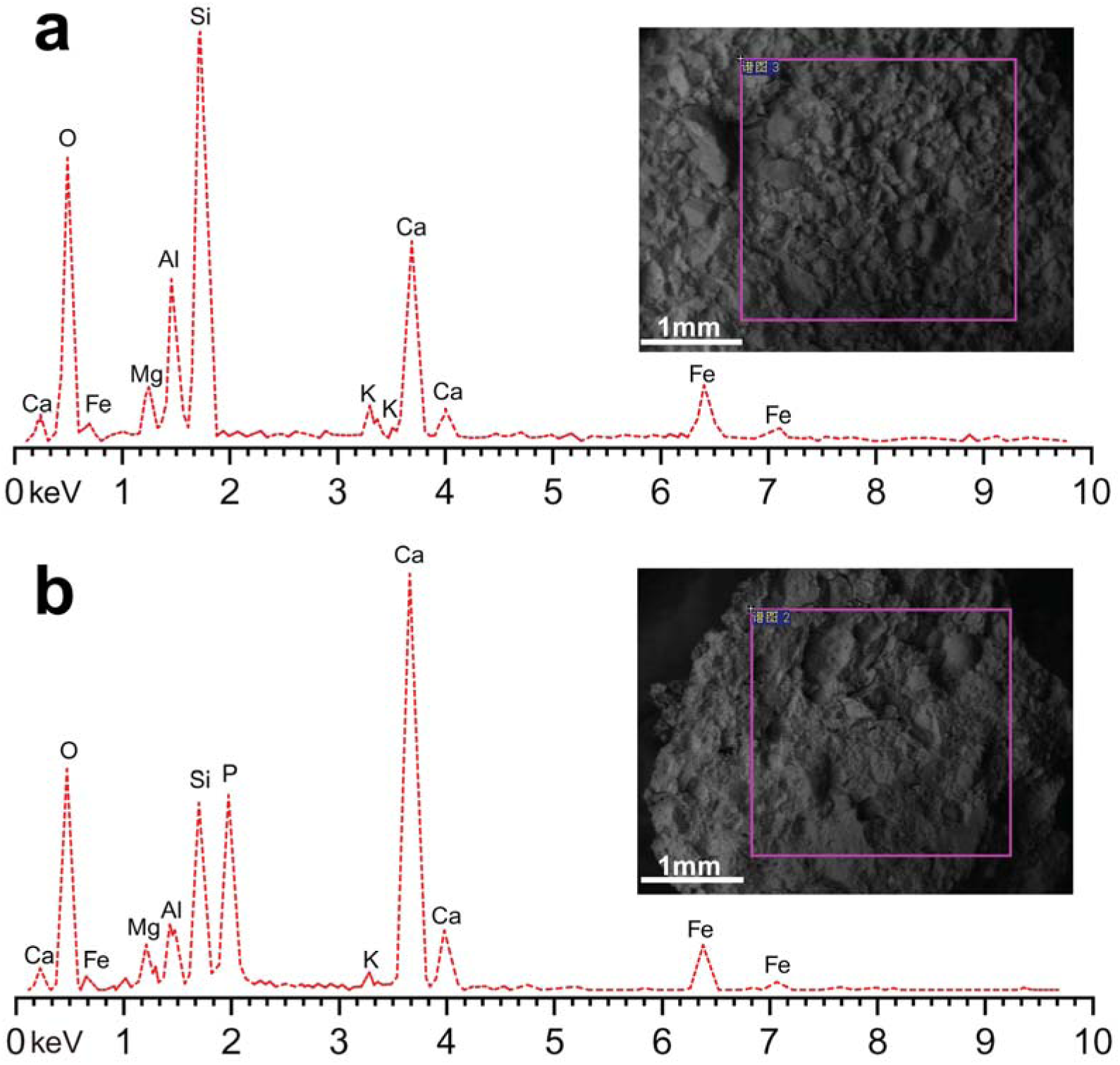
EDS results for the yellowish conglomerate enclosed by the partial gastralia ribcage. **a,** EDS spectra derived from the host sandstone show the high levels of O, Si, Al, and Ca, which are commonly enriched in the Earth’s crustal rocks; **b,** EDS spectra derived from the conglomerate highlighting high concentrations of Ca and P, indicating a carcass origin of these elements.

**Figure 3.**
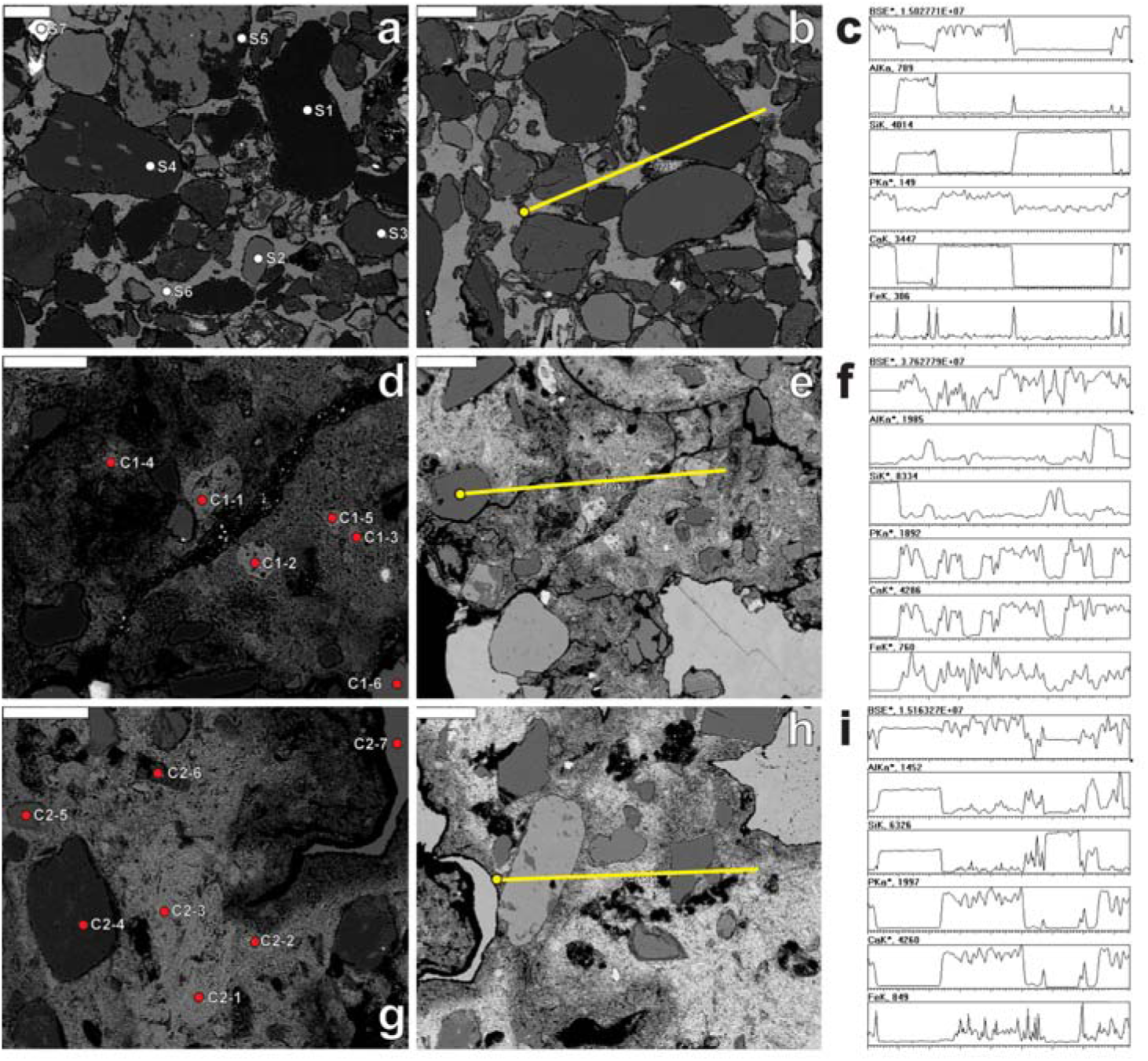
Ground sections and EMP results highlighting the differences in microstructures, chemical components, and roundness of the minerals cemented in the host sandstone and conglomerate samples. **a-b,** Ground sections of the host sandstone showing the cemented mineral particles. The white spots mark the positions where chemical components were detected (Extended Data Table 1), with the changes in chemical components along the testing tract (yellow line) are shown in **c**; **d** and **g,** Ground sections of the conglomerate samples (1 and 2, respectively) showing the massive apatite with tiny granular structures, indicating the presence of hollow microspheres. The red spots mark the positions where chemical components were detected (Extended Data Table 1), with the changes in chemical components along the testing tract (yellow lines) are shown in **f** and **i**, respectively. **Scale bar**=200µm. **Abbreviations: S,** sandstone; **C,** conglomerate.

Scanning electron microscopy (SEM) analysis revealed tightly packed hollow microspheres, each with a diameter of less than 2µm, within the conglomerate where apatite was present (Fig. 4 a-c). These microspheres, which differ from the rod-like phosphatized bacteria (Fig. 4g), are identical in size, shape, and chemical composition to those found in the intestinal remnants of *Scipionyx*[2], *Mirischia*[30], and the coprolites of the Late Triassic archosaur *Smok wawelski*[9]. This similarity suggests that the original tissues were largely replaced by phosphatized microbes during fossilization (microbial microfabric[13]). Film-like apatite clusters were observed nearby (Fig. 4 d-f), indicating a heterogeneous calcium phosphate-enriched substrate (Fig. 4 h-i). Since little mineral particles from the surrounding sandstone were found within these film-like apatite clusters, it is likely that they formed through microbe-mediated phosphatization in the relatively isolated microenvironment without the direct mineralization of the microbes themselves (e.g., intermediate microfabric[13, 31]). Both types of apatite differ from the crystalline apatite typically found in igneous and metamorphic rocks[32]. In addition, phosphatized filaments found atop the apatite substrates (Fig. 5 a-i) likely represents phosphatized soft tissues. Debris and imprints with longitudinal stripes are found within the apatite matrix, pointing to the presence of hard tissues (Fig. 5 k-l). These are the most direct indicators of intestinal food remains. The heterogeneous composition of the conglomerate suggests that it is made up of materials with varying degrees of resistance to decay, rather than being derived from the carcass of the host animal.

**Figure 4.**
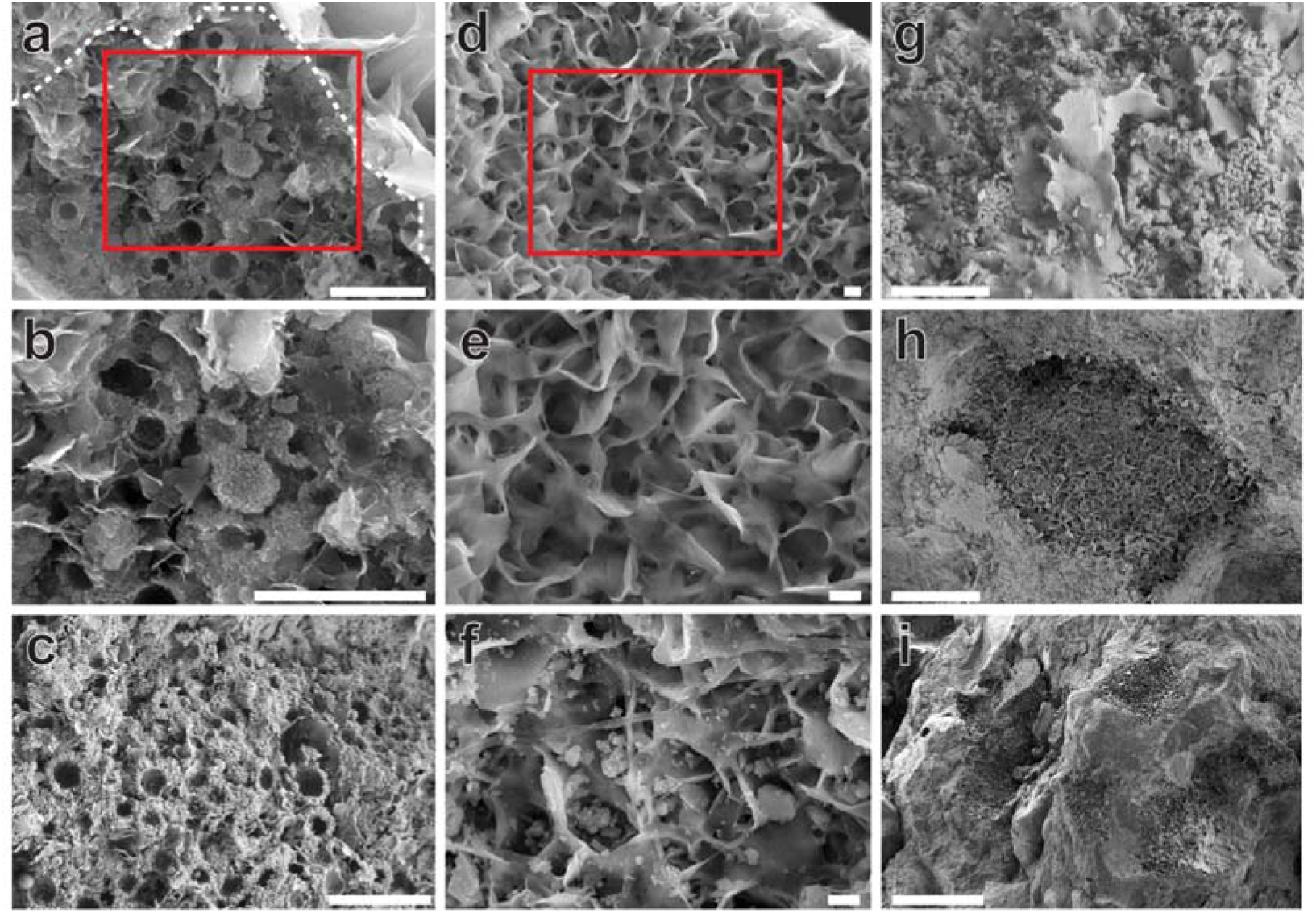
SEM images showing the phosphatized hollow microspheres (a-c), apatite crystals (d-f), and phosphatized microbes (g), illustrating the heterogeneous nature of the conglomerate (h-i). Notably, the phosphatized hollow microspheres, with diameters no less than 2µm (**a-c**), resemble the pseudomorphed soft tissues within the intestinal contents of *Scipionyx samniticus*[2]. The apatite crystals (**d-f**) represent a type of intermediate microfabric that are previously unknown. The red boxes in **a** and **d** highlight the areas that are magnified in **b** and **e**, respectively. **Scale bars** are set at 5µm for **a-f**, 1µm for **g**, 100µm for **h**, and 250µm for **i**.

**Figure 5.**
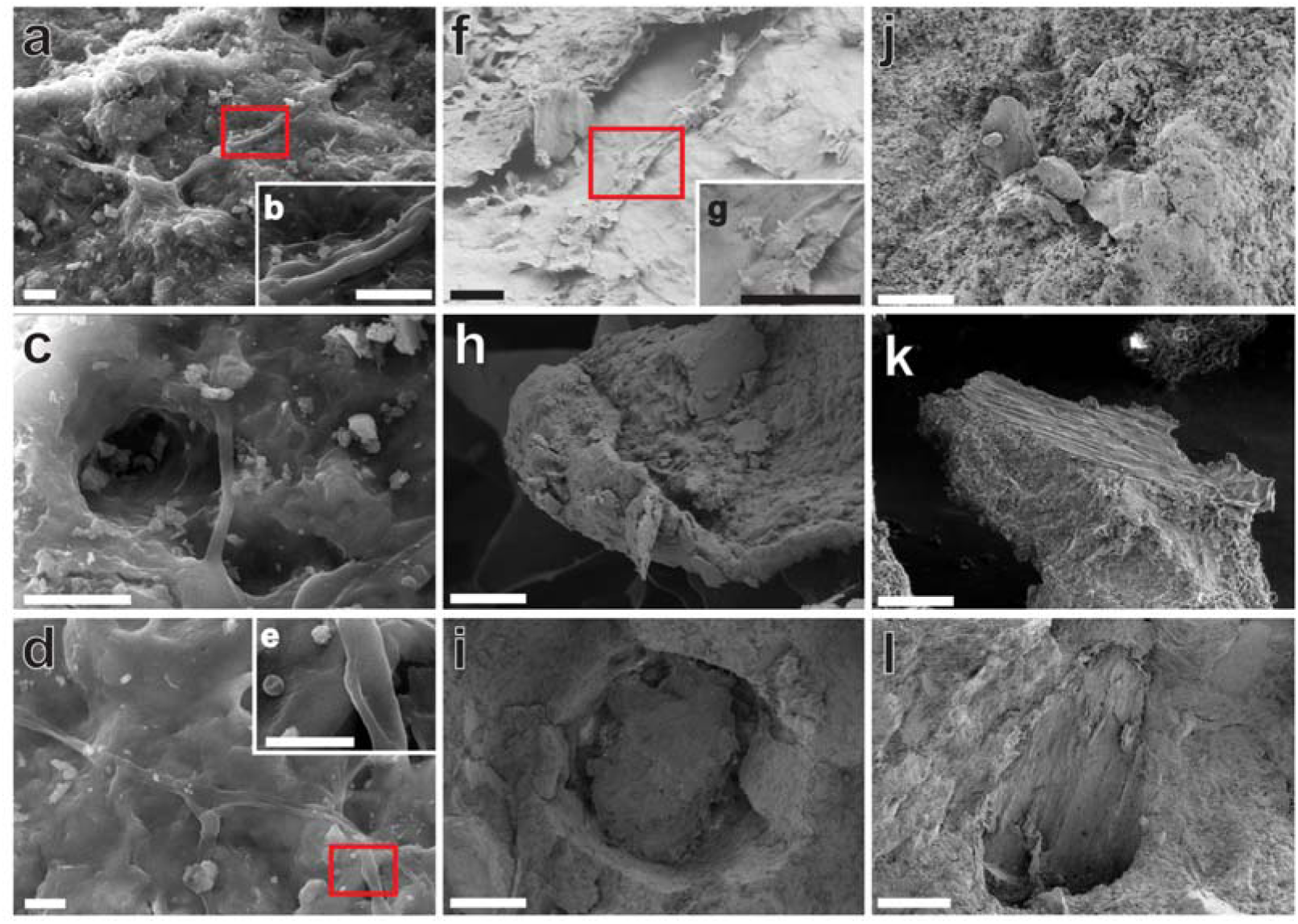
SEM images showing the phosphatized soft tissues (a-i) and debris of hard tissues (j-l) within the intestinal contents. Notably, the debris and imprints of hard tissues are marked by longitudinal stripes (**j-l**). The red boxes in **a, d** and **f** highlight the areas that are magnified in **b, e** and **g**, respectively. **Scale bars** are set at 5µm for **a-e** and **h**, 3µm for **f-g**, 50µm for **i**, 25µm for **j-k**, and 100µm for **l**.

To further confirm the sources of the food remains, Raman spectrum (RS) analysis was conducted on both the hard tissue debris and the surrounding sandstone. Featured peaks characteristic of fully fossilized bony tissue were present in the remains samples but absent in the surrounding sandstone (Fig. 6, Table 1). Previous study showed that these peaks remain stable across different localities but are closely correlated with the degree of fossilization, suggesting that diagenesis, along with the presence of bony substrate, are crucial in generating these feature peaks[33]. Given that no skeletal elements are expected within the gastralia ribcage, the result suggests both the phosphatized soft tissues and debris of hard tissues were derived from dissolved bony substrates of prey[2, 9, 34]. No peaks indicative of plant remains or insect chitin were detected (Extended Data Fig. 2), indicating that plants and insects were not part of the last meal of *Bannykus*.

**Figure 6.**
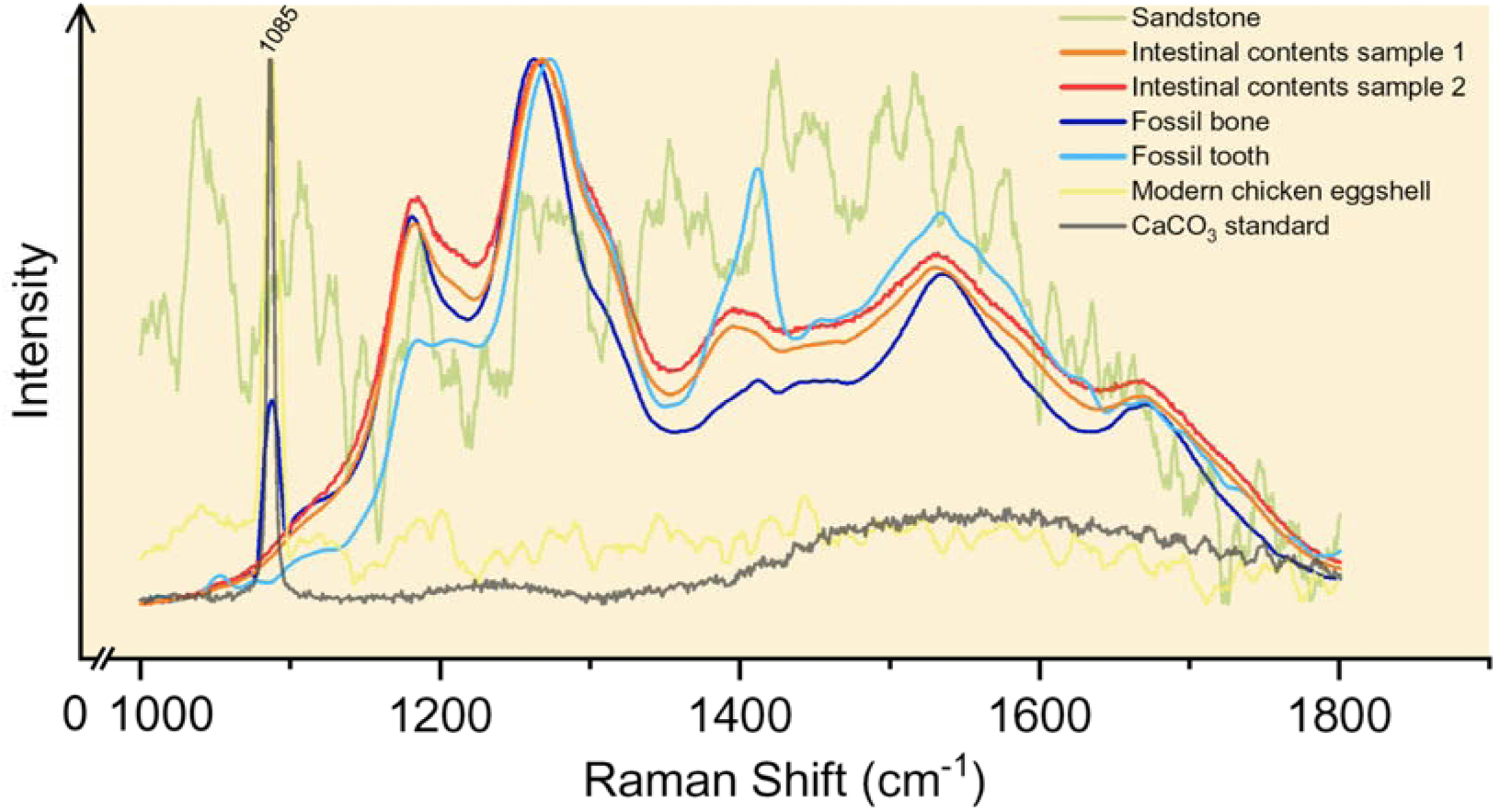
Raman spectrum results illustrating feature peaks indicative of bony substrates of the intestinal content. Raman spectra are collected with 785 nm excitation wavelength, which is common in fossil materials [33]. In addition, the wavelength does not alter the number or positions of the feature peaks. Fossil bone and tooth from geographically and chronologically disparate localities were included to eliminate potential biases in the Raman signals of the intestinal content samples. An eggshell and calcium carbonate standard were included to assess the possibility of egg formation in the abdominal cavity. Notably, a peak at approximately 1085 cm^−1^, corresponding to the symmetric stretching vibration (ν□) of the carbonate ion (CO^2−^) was observed in the eggshell, calcium carbonate standard, and fossil bone. In contrast, this peak was absent in the intestinal content samples, reinforcing the hypothesis that the stomach contents are primarily composed of apatite, with minimal calcium carbonate, thereby ruling out contamination from eggshell material. Furthermore, a series of shared characteristic peaks between 1100 cm^−1^ and 1700 cm^−1^ observed in both the intestinal contents and the fossil bone/tooth may reflect carbonate-related signals in the well-fossilized bone tissue. Despite variations in burial locations and geological periods, and given that tooth enamel is generally more resistant to diagenetic processes than bone, the similarity of these peaks suggests potential diagenetic alterations.

**Table 1.** Correlation analysis of Raman spectrum further strengthens the evidence for the biological apatite origin of intestinal contents.

| Pearson's correlation coefficient | Standstone | Intestinal contents sample 1 | Intestinal contents sample 2 | Fossil bone | Fossil tooth | Modern chicken eggshell | CaCO<sub>3</sub> standard |
| --- | --- | --- | --- | --- | --- | --- | --- |
| Standstone | 1 | 0.40194 | 0.3954 | 0.3862 | 0.49802 | 0.31412 | 0.07783 |
| Intestinal contents sample 1 | 0.40194 | 1 | 0.99866 | 0.9734 | 0.94273 | -0.0902 | -0.0344 |
| Intestinal contents sample 2 | 0.3954 | 0.99866 | 1 | 0.9701 | 0.93639 | -0.0934 | -0.0365 |
| Fossil bone | 0.38619 | 0.97343 | 0.97013 | 1 | 0.88314 | 0.04347 | 0.04622 |
| Fossil tooth | 0.49802 | 0.94273 | 0.93639 | 0.8831 | 1 | -0.0813 | 0.06868 |
| Modern chicken eggshell | 0.31412 | -0.0902 | -0.0934 | 0.0435 | -0.0813 | 1 | 0.58748 |
| CaCO<sub>3</sub> standard | 0.07783 | -0.0344 | -0.0365 | 0.0462 | 0.06868 | 0.58748 | 1 |

The Pearson correlation coefficient measures the linear relationship between two variables, ranging from -1 to 1:
1 indicates a perfect positive linear relationship: as one variable increases, the other also increases proportionally;
0 indicates no linear relationship between the variables;
-1 indicates a perfect negative linear relationship: as one variable increases, the other decreases proportionally.

## Discussion

The potential ecological niche, including the possible diet and feeding strategies of alvarezsaurians, remains poorly understood due to the lack of direct dietary evidence. The incompleteness of cranial elements and the complex trait evolution within this clade further complicate this issue[27]. Among alvarezsaurians, relatively complete cranial elements are only known from *Haplocheirus* and *Shuvuuia*. A preliminary finite element analysis (FEA) shows that *Haplocheirus* had a jaw adductor muscle force comparable to that of the tyrannosaurid *Dilong*[35], and a bite force over ten times greater than that of *Shuvuuia* (Extended Data Fig. 1, Extended Data Tables 1-4). This suggests that early- and late-diverging alvarezsaurians may have occupied different ecological niches and employed varying feeding strategies, consistent with their morphological differences.

For alvarezsaurians lacking available jaw morphologies, dietary inference largely relies on alternative evidence. Although little is known about the jaw and teeth morphology of *Bannykus*, certain features such as the poorly defined supratemporal fossa, the absence of a mandibular infracondylar fossa, and the undeveloped surangular crest suggest that its jaw may resemble those of late-diverging parvicursorines like *Shuvuuia* rather than *Haplocheirus*[27]. This implies that the diet and feeding strategies of *Bannyskus* may have been more similar to *Shuvuuia* than *Haplocheirus*. However, the discovery of bony tissues within the intestinal contents suggests that *Bannykus* may have retained traces of its ancestral dietary habits. This finding reinforces the intermediate status of *Bannykus*, where morphological changes, such as the attenuating of the cranial elements and the initiation reduction of manual digits, occurred before shifts in diet (Fig. 7). This makes an in-depth analysis of its intestinal contents especially valuable for understanding *Bannykus*’s feeding strategies.

**Figure 7.**
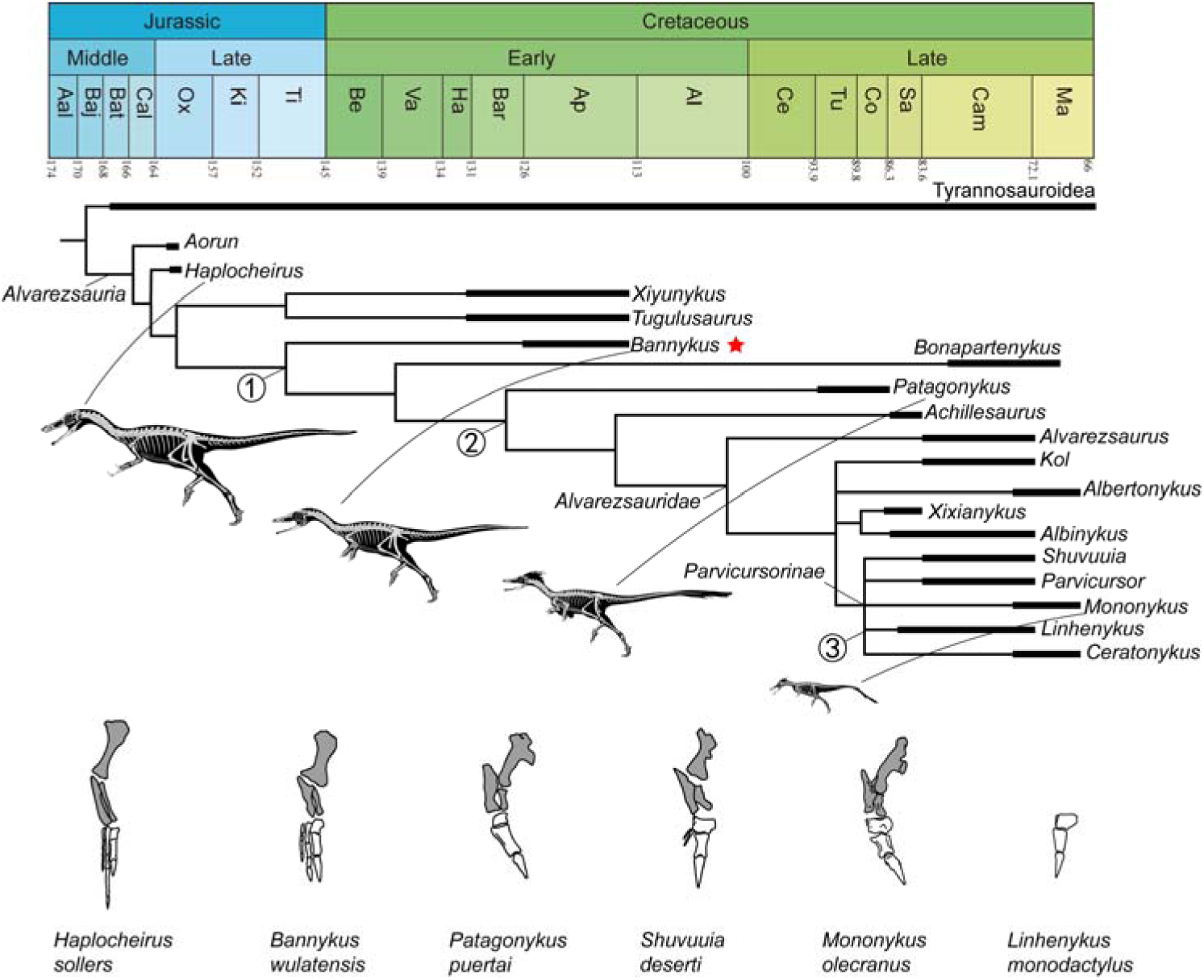
Time-calibrated simplified phylogeny showing the interrelationships of alvarezsaurians, along with the accompanying decreases in body size and the reduction of manual digits (not to scale). Three major events of manual digital reduction occurred during the evolution of alvarezsaurians, with manual digit III being reduced first ①, followed by further reduction of the lateral digits (III and IV) ②. Only manual digit I is present in *Linhenykus*, suggesting the stylopod and zeugopod are further reduced in parvicursorines, leaving the manual digit II the only developed forelimb element. Our results indicate that the ancestral carnivorous diet remained when manual digit III began to reduce, and that body miniaturization, manual digit reductions and the possible dietary shift were not synchronized. The alvarezsaurian phylogeny[27], skeletal silhouettes[67] and schematic drawings of forelimbs[68] were respectively modified from previous works.

Our analysis reveals a notable enrichment of calcium phosphate in the intestinal contents of *Bannykus*. The phosphorus content in the Earth’s crustal rocks is typically around 0.1%[36], and it is significantly lower in soft tissues compared to bones. A similar high concentration of calcium phosphate is also observed in the gastrointestinal contents and coprolites of several bone-consuming animals[2, 9, 34, 37]. In contrast, the insect exoskeletons, made of chitin[38], are primarily composed of carbon and contain almost no phosphorus (Extended Data Fig. 2) similar to plant materials[38, 39]. These findings strongly suggest that *Bannykus* has a carnivorous diet.

Although the pseudomorphed soft and hard tissues provide direct indicator of diet, identifying the source of prey remains challenging due to various biological and non-biological factors. Angular bones fragments found in the posterior digestive tract might indicate bone-crushing behavior, a short gut retention period, limited chemical digestion, or a combination of these factors[8, 29], while identifying more complete and anatomically connected structures may indicate whole-prey swallowing together with minimum digestion[3, 4]. Typically, identifiable digestive remains come from the anterior half of the digestive tract[3, 4, 6], where digestion is just beginning and food is less degraded[3, 4]. In contrast, material from the duodenum-to-rectum often lack diagnostic features due to effect of mechanical and chemical digestion[2]. Physiological factors, such as stomach acid strength, digestive enzymes, digestive tract structure, and gut retention time, affect the identification of these remains[29, 31, 34]. Non-physiological factors, including meal frequency[2, 29], microbe activities[13, 31, 34, 40], sediment pH[31, 34, 40, 41], and taphonomic conditions[13], also play significantly roles in the preservation and identification of dietary remains. The composition of stomach contents is often be inferred by identifying associated animals from the same locality. For example, digested enantiornithines and lizards have been identified in the gastrointestinal remains of *Microraptor*[3, 4]. However, the absence of associated fossils at the locality where *Bannykus* was found limits the estimation of its diet and the use of indirect methods, such as isotopic geochemistry.

Form and function are related, and the intestinal contents or coprolites can reveal more about their feeding strategies and digestive physiology than specific dietary components. For instance, crocodiles swallow large chunks of food whole and rely on strong chemical digestion[42, 43], leading to feces that typically contain no bone fragments[42]. Similarly, snakes regurgitate indigestible materials[44], while birds have developed a two-part stomach that grinds food and aids in the regurgitation of indigestible materials[45], resulting in little to no indigestible material in their feces[46]. Osteophagous animals, like hyenas, have developed powerful bite force to crush bones[47], leading to feces with highly digested bone fragments[47].

Although reptiles are generally less effective at oral food processing than mammals (e.g., hyena), theropods like *Allosaurus* and *Tyrannosaurus* are thought to have had bone-crushing abilities[9, 48]. These theropods likely swallowed large prey while cracking some bones, as opposed to swallowing prey whole like snakes. Pellet egestion, believed to have first appeared in *Anchiornis*[6, 46], likely helped reduce gut retention time and increases digestive efficiency in paravians including birds. Other carnivorous non-avialan theropods, including *Microraptor*, are thought to have swallowed prey whole and excreted most indigestible matters through their feces[46]. These raptors possessed powerful chemical digestion, with a gastric pH that is typically lower than 2[49]. The high concentration of calcium phosphate of *Bannykus*, along with the appearance of hard tissues, suggests that most hard tissues including bony material was fully digested before reaching the duodenum-to-rectum region[49]. This digestive process may be more similar to that of crocodiles than to most other non-avialan theropods, making *Bannykus* unique among established carnivorous theropods.

While evidence suggests that bony tissues were part of *Bannykus*’s last meal, it does not exclude the possibility of other food resources. The trend of utilizing a diverse range of trophic resources is well-documented in both avialan and non-avialan theropod dinosaurs[1, 15, 46]. An extreme example is seen during the ontogenetic development of *Limusaurus*, where young juveniles possessing teeth and gastroliths were shown to be omnivorous, while older, edentulous individuals were herbivorous[16]. Similarly, an egested pellet associated with *Iteravis* contained bone fragments, carbonized remains, and a small cluster of gastroliths[46], suggesting that the presence of gastroliths does not necessarily indicate an herbivorous diet[1, 46]. Reports of opportunistic feeding behaviors in *Microraptor*[1, 4, 50, 51] and *Anchiornis*[6], where various food resources including aquatic, terrestrial, and arboreal animals were found within the stomach contents, further illustrate this diversity. Although current lines of evidence are insufficient to confirm whether *Bannykus* was a strict carnivore, scavenger, or opportunist; however, its long legs suggest adaptations for active hunting, consistent with a predatory lifestyle.

The ability to digest chitinous substrates is deeply rooted in theropod ancestors, as evidenced by chitinous insect cuticles found in coprolites potentially belonging to the Triassic dinosauriform *Silesaurus opolensis*[52, 53]. Despite current lines of evidence support a carnivorous diet for *Bannykus*, it does not entirely rule out the possibility of an insectivorous diet suggested for late-diverging parvicursorines. If a dietary shift to insectivore occurred, it likely took place later in alvarezsaurian evolution, probably coinciding with a reduction in body size[26].

## METHODS

### Scanning electron microscopy coupled with energy-dispersive X-ray spectroscopy

Samples taken from the conglomerate and host sand stone were analyzed using a Hitachi S—3700N (Institute of Vertebrate Paleontology and Paleoanthropology, Chinese Academy of Sciences) and 4800 (East China Normal University) scanning electron microscope (SEM) equipped with the EDS detector, and was operated at an accelerated voltage of 25 kV, working distance of 8.5~8.7mm, and live time of 4 seconds.

### Electron microprobe

Quantitative constituent elements analysis of samples taken from the conglomerate and host sand stone were analyzed using electron probe microanalyzer JAX-8100 at the Institute of Geology, Chinese Academy of Geological Sciences. The routine microprobe analyses were performed in the condition of 15 kV, with a beam current of 10 nA and a beam spot of 1~5µm.

### Finite element analysis

The biomechanics of alvarezsaurian mandibles were studied using finite element analysis. The early-diverging alvarezsaurian *Haplocheirus sollers* and the late-diverging alvarezsaurid *Shuvuuia deserti* were included in the study. The digital models of the mandibles of *Haplocheirus* (IVPP V14988) and *Shuvuuia* (MGI 100/977) were obtained, which were generated via segmentation of computed tomography (CT) scan data. Due to limitation of the resolution of the CT data, cortical and cancellous bone were not distinguished in the models. To minimize the influence of postmortem deformation on biomechanical analysis, we performed retrodeformation of the mandibles in the three-dimensional modelling software Blender (version 2.79b; www.blender.org). This process includes filling in breaks and cracks, replacing missing elements by reflecting bilateral elements, duplicating serially repeated elements, modifying from closely related taxa, repositioning or disarticulated elements, and correcting plastic deformation of elements (e.g., [54–56]). Anatomical details not captured in the CT scan data were adjusted with reference to firsthand examination of the original and referred specimens (MGI 100/1001 for *Shuvuuia*) and relevant osteological descriptions.

The setup for finite element analysis was conducted in the software *Hypermesh* (version 11, Altair Engineering Inc.). The mandible models of *Haplocheirus* and *Shuvuuia* underwent solid meshing to produce meshes consisting of tetrahedral elements (4616244 and 1300082 elements respectively). The material properties of the mandibles of extant alligators (Young’s modulus (E) = 20.49 GPa, Poisson’s ratio (□) = 0.4) were used for the alvarezsaurians [57], as in other studies on non-avialan theropod dinosaurs (e.g., [58, 59]). The material is isotropic and homogeneous.

Loading conditions simulating biting were set up to test the mechanical response of the mandible under feeding. The origin and insertion sites of jaw adductor muscles were identified using a phylogenetic bracketing approach based on extant archosaurs [60], with reference to the reconstructions of closely related theropod clades [55, 56, 61] (Extended Data Table 3). The forces generated by the adductor muscles were calculated using dry skull method [62]. Due to the unavailability of digital models of the cranium, three-dimensional volumetric reconstruction of adductor muscles was not feasible in our study. While the dry skull method has been observed to underestimate/overestimate muscle force in mammals [63] and potentially influence the absolute values, it remains the optimal alternative herein considering the available materials. The primary objective of the biomechanical analysis is to compare the basal and more derived conditions in alvarezsaurians. As the method of force calculation remains consistent across both models, this approach is likely to yield reliable comparative results for the identification of potential changes in feeding mechanics throughout the evolution of alvarezsaurians.

The adductor muscle insertion sites were first mapped on the mandible models. Using published and firsthand photos of the skull of the species, muscle origins were identified with the adductor muscles mapped in lateral view. Three functional muscle groups were differentiated: pterygoid muscle group (mPTd, mPTv), quadrate muscle group (mPSTp, mAMP), and temporal muscle group (mPSTs, mAMEP, mAMEM, mAMES), as similarly implemented in [48, 64, 65]. The physiological cross-sectional areas of the muscle groups were estimated in consideration of the dimension of the skull hard tissue bounding the jaw adductor muscles[64], based on published and firsthand photographs of the original and referred specimens. To estimate the physiological cross-sectional area of pterygoid and quadrate muscle groups, the anterior and posterior extents of the muscles (based on the muscle reconstructions in lateral view) were first mapped on the ventral view of the skull. The ventral area, constrained by the anteroposterior boarders and the space available mediolaterally between jugal/quadratojugal and medial surface of the hemimandible, was measured. Similarly, the dorsal area of the temporal fossa was measured to estimate the force generated by the temporal muscle group. All of the measurements were conducted in the software ImageJ [66]. The muscle force generated by the muscle group(s) is calculated as the product of the measured physiological cross-sectional area and an isometric muscle stress value (30 N/cm^2^) [62]. To obtain the force magnitude of each individual muscle for setting up the loading conditions of finite element analysis, the estimated force of a muscle group was divided by the total number of individual muscles, assuming each muscle contributes the same ratio of force (Extended Data Table 4). The direction of the force vector was determined using the mandible models and photos of the skull of the species.

Three bilateral bite scenarios were set up: at the first tooth, middle of the toothrow (17th for *Haplocheirus*; 20th for *Shuvuuia*), and the last tooth. The mandible models were constrained at 16 nodes at the mandibular joints from x-, y- and z-direction movement. Two additional constraints were placed at the bite points for the respective bilateral bite scenario.

Finite element analysis was performed in Abaqus (version 6.141, Dassault Systèmes Simulia Corp.). Contour plots of von Mises stress and were obtained. Several parameters were used to evaluate and compare the feeding mechanics of the mandible models. The average stress experienced by the mandible was calculated, which was defined as the total stress experienced by the mandible divided by the number of elements. The reaction forces at the bite points were recorded for bite force estimations and the calculation of mechanical advantage of the biting system. Mechanical advantage is defined as the percentage of the total input adductor muscle forces converted to bite force (Extended Data Table 1~4).

### Raman spectroscopy

Raman spectroscopy was conducted using a Renishaw inVia-Reflex confocal Raman spectrometer, equipped with a research-grade Leica microscope and a 785 nm semiconductor laser. The instrument was operated with a 50× objective. The spectra were acquired over a wavenumber range of 100 to 3200 cm^1^. The laser power on the sample was kept below 5% of its maximum power, with an exposure time of 10 seconds per spectrum.

Seven samples were analyzed: two intestinal contents samples prepared from inside the ribcage, the surrounding sandstone, a piece of bone fragment of an unnamed fossil lizard collected from Inner Mongolia, China, a piece of tooth enamel of an unnamed sauropod recovered from Wucaiwan, Xinjiang, China, a piece of eggshell from a modern chicken and some powder of the calcium carbonate standard.

Data were smoothed and baseline-corrected using Renishaw’s WiRE 5.1 software. The spectral data ranging from 1000 to 1800 cm^1^ were imported into OriginPro 2019b software for further analysis and plotting. The Pearson correlation coefficients between the normalized intensity data of each sample were then calculated to evaluate the similarities between the samples.

The Pearson correlation coefficients between the different samples included in the Raman spectrum analysis are presented. The coefficients range between −1 and 1, where values closer to 1 indicate a strong positive linear relationship.

## Supplementary Information

is available in the online version of the paper.

## Acknowledgments

We thank L. Xiang, T. Yu, and X. Ding (IVPP) for preparing the specimen, and H. Zang (IVPP) for photographing the specimen. We also appreciate the invaluable help of L.T. (Long Hao Institute of Geology and Paleontology) in collecting this specimen. Just as this manuscript was being completed, we were deeply saddened to learn of the passing of Professor Tan Lin. This article is dedicated to Professor Tan Lin for his great contributions to vertebrate paleontology in Inner Mongolia. S.W. was supported by the Zijiang Program for Talented Scholars at East China Normal University and the Shanghai Pujiang Program (23PJ1402300); X.X. was supported by the National Natural Science Foundation of China (Grant No. 42288201) and the Yunnan Revitalization Talent Support Program (202305AB350006); W.M.’s participation in this study was supported by the Deep Time Peter Buck Postdoctoral Fellowship from the National Museum of Natural History, Smithsonian Institution.

## Author Contributions

S.W. and X.X. conceived the project; S.W. and N.D. performed the SEM, EDS, and EMP analyses, W.M. performed the finite element analysis, W.Y., N.D. and T.Z. performed the RS analysis, J.C. help reconstruct the 3D model of *Shuvuuia*, X.X. led the fieldwork and collected the specimens; and S.W. wrote the manuscript.

## Author Information

Correspondence and requests for materials should be addressed to S.W. regarding the dietary analysis, or X.X. regarding the specimen.

